# BENDER DB: a database of protein binding sites across neglected disease proteomes

**DOI:** 10.1101/2025.02.18.638856

**Authors:** Vinícius A. Paiva, Douglas E. V. Pires, Gustavo C. Bressan, Sandro C. Izidoro, Sabrina A. Silveira

## Abstract

Identifying binding sites is crucial for expanding our knowledge of various biological processes, supporting drug discovery, and repositioning strategies, particularly in the early research phases of target hopping. This is especially important for neglected diseases, which predominantly impact vulnerable populations in developing regions. To address this, we created BENDER DB, a database designed to map predicted protein binding sites within the proteomes of pathogens associated with neglected diseases. Utilizing AlphaFold-predicted structures, BENDER DB integrates results from five leading binding site prediction tools, resulting in over one million binding sites across over 100,000 proteins from 10 different proteomes. BENDER DB offers unique features, such as integrating multiple predictors, interactive visualization tools, and comprehensive graphical representations, allowing for a detailed visual comparative analysis of different prediction methods and the predicted binding sites. We combined the computational approaches to offer a consensus of binding site predictions, leveraging the strengths of multiple techniques. Additionally, we introduce BENDER AI, a meta-predictor that combines the outputs of the predictors to provide a unified binding site classification. BENDER AI outperformed all individual methods in 4 out of 5 evaluated metrics. Specifically, it achieved an MCC of 0.64 and an AUC of 0.89, surpassing the highest MCC of 0.63 and the highest AUC of 0.84 obtained by individual predictors, demonstrating strong performance and effectiveness in supporting binding site prediction. By consolidating these results, BENDER DB enables detailed comparative analyses of prediction methods and integrates interactive visualization tools to provide users with an intuitive platform for binding site exploration. The database aims to accelerate drug discovery efforts, particularly in underdeveloped regions, by providing a robust and detailed resource for studying protein binding sites. BENDER DB is available at https://benderdb.ufv.br.

## 1. Introduction

The UniProt database, a comprehensive repository for protein sequences and their functions, currently contains over 248 million annotated protein sequences. Of these, 572,214 entries are reviewed and curated in the Swiss-Prot section, providing high-quality, expert-verified annotations. The remaining 248,266,673 entries in TrEMBL are unreviewed and translated directly from nucleotide sequences [1]. Despite this extensive dataset, the number of proteins experimentally characterized through methods like X-ray crystallography, Nuclear Magnetic Resonance (NMR) spectroscopy, and Cryo-Electron Microscopy (Cryo-EM) remains limited due to the high costs and time-intensive nature of these processes [2, 3]. Additionally, certain protein families present their challenges. For example, membrane proteins, such as G protein-coupled receptors (GPCRs), which play a fundamental role in various biological functions and are common targets in the development of new drugs, have their annotation hindered by their low levels of native expression and low intrinsic stability [4, 5, 6, 7].

Identifying binding sites is essential to broaden our understanding of different biological processes. It can support drug discovery and drug repositioning strategies, especially in the early stages of research, such as target hopping [8, 9, 10, 11]. Computational methods have been proposed to search for these valuable regions, offering cost and time savings compared to experimental procedures. These methods, which typically employ approaches involving Machine Learning and Deep Learning techniques, increasingly utilize protein features such as physicochemical properties, binding affinities, and traditional information on structure and sequence [12, 13, 14]. Most predictors depend on annotated data, although some new models have been developed using protein language and unsupervised learning models [15, 16, 17].

Both structural and sequence characteristics are critical in binding site prediction [18]. While some approaches focus on sequence-based features, such as local sequence motifs or conserved residues, others incorporate structural information to enhance prediction accuracy [12, 19]. Sequence-based techniques often analyze segments of protein sequences to predict binding residues, as seen in approaches like Fuzzy Cognitive Maps (FCM), which evaluate local characteristics to predict DNA-binding residues [20]. On the other hand, structure-based methods examine the 3D conformation of proteins, leveraging properties like surface geometry, electrostatic potential, and hydrophobicity to identify potential ligand-binding pockets [21, 22].

Numerical and computational methods form the foundation of modern binding site predictions, including those used in BENDER DB. These methodologies have been applied successfully in various areas of medical research, including drug discovery, disease modeling, and therapeutic optimization [23, 24]. An example of the application of such numerical techniques is the use of fractional-order dynamics in medical models, which offers a deeper understanding of biological processes through mathematical simulations. Studies exploring fractional-order dynamics [25, 26] demonstrate how advanced numerical approaches can contribute to drug discovery by modeling complex systems.

Binding site prediction methods are crucial in addressing diseases with limited therapeutic options [27, 28]. For instance, identifying potential drug targets is vital in neglected diseases, which predominantly affect vulnerable populations in underdeveloped and developing regions of Latin America, Asia, and Africa. These diseases, characterized by poor health indicators and minimal investments in research and control, cause significant morbidity and mortality, with an estimated 200,000 deaths annually [29, 30]. Therefore, accurate prediction of binding sites can accelerate the identification of viable drug targets, ultimately aiding in developing effective treatments for neglected diseases.

A database that catalogs protein binding sites would be a valuable tool in studying neglected diseases. A user-friendly resource could democratize access to critical data, allowing researchers to overcome the barriers posed by the need for computational expertise and the scalability challenges of many binding site prediction methods. By compiling results from various predictors, such a database would overcome the limitations of individual techniques, providing a consensus that enhances the reliability and accuracy of binding site identification. This is particularly important in neglected diseases, where identifying potential drug targets is crucial for developing effective therapies for vulnerable populations.

In this context, we developed BENDER DB (a database of protein Binding sitEs across Neglected DiseasE pRoteomes) to comprehensively map putative/predicted protein binding sites within the proteomes of neglected disease pathogens. The primary goal of BENDER DB is to identify and catalog protein binding sites using five different binding site predictors, thereby creating a robust and detailed resource. This database features over 1 million binding sites across more than 100,000 proteins in 10 different proteomes, highlighting commonalities across predictors and leveraging visualizations to aid researchers in pinpointing promising targets and designing practical drug discovery experiments.

Representative examples of databases related to protein-ligand binding sites are: (i) BindingDB [31], which is a publicly accessible resource that contains experimental protein-small molecule interaction data. Its entries are sourced from scientific literature and US patents; (ii) Binding MOAD [32], which provides a subset of Protein Data Bank (PDB) [33] entries, representing high-quality x-ray crystal structures solved with biologically relevant ligands bound; (iii) BioLip [34], which is a semi-manually curated database that focuses on high-quality, biologically relevant protein-ligand interactions from the PDB. Regarding databases related to binding sites using automatically predicted structures, we can mention: (iv) HProteome-BSite [35], which catalogs protein binding sites within the human proteome using structures predicted by AlphaFold [36, 37]. It offers a comprehensive database for studying protein-ligand interactions, focusing on functional sites in human proteins.; (v) CavitySpace [38], which identifies and catalogs potential binding pockets in protein structures obtained from the AlphaFold database [39]. Its data includes detailed information on cavity sizes, shapes, and locations.; (vi) AlphaFill [40] enhances AlphaFold structure models by integrating ligand information from experimentally determined PDB structures. The database provides annotations of ligand binding sites within these extended models.

While databases like those provide valuable information on protein binding sites, BENDER DB offers unique features that set it apart. It takes as input structures predicted by AlphaFold, comprising the whole proteome of neglected diseases. Our tool integrates binding site prediction results from five state-of-the-art predictors, consolidating them in a single location and highlighting their similarities and differences. It catalogs proteins on a proteomic scale within the context of neglected diseases, providing a comprehensive mapping essential for identifying promising drug targets. Additionally, BENDER DB includes visual interactive tools and graphical resources that enhance the study of binding sites. It is a more versatile and specialized resource for researchers focused on neglected diseases.

A key feature BENDER DB offers is the ability to highlight how the binding site predictions of the various methods converge. Upon searching for the protein in the database, users are presented with a results page summarizing the information from several binding site predictors. This page lets users explore each prediction model individually, enabling a detailed and customized analysis. Graphical representations of the protein binding sites are displayed through a molecular visualization, including a consensus tab, which aggregates all prediction results from various methods into a single representation. Users can also analyze different sets and intersections between the predictor’s results using an UpSet plot [41], a graphical visualization designed to analyze set intersections. We also developed BENDER AI, an artificial intelligence model designed to identify binding sites in proteins by integrating the outputs of all the predictors included in BENDER DB. With these comprehensive tools, we offer a robust and versatile platform for analyzing proteins and their binding sites, serving as a valuable resource for researchers.

## 2. Related works

Several databases have been developed to catalog protein binding sites, providing critical resources for drug discovery and structural biology research. These databases serve as valuable tools in identifying potential therapeutic targets by cataloging experimentally verified or computationally predicted binding sites in protein structures.

Among the experimental resources, BindingDB [31] is a comprehensive database that collects protein-small molecule interaction data from scientific literature and patents. This wealth of experimental data helps understand protein-ligand interactions in known structures. Still, it is limited to available experimental data and does not provide predictions for proteins lacking such information. Similarly, Binding MOAD [32] focuses on high-quality, x-ray crystallographic data from the Protein Data Bank (PDB), representing biologically relevant ligand-bound structures. While these resources are invaluable for studying proteins with experimentally solved structures, their reliance on empirical data limits their coverage, particularly for proteins from neglected disease pathogens, where structural information is often scarce.

The BioLip database [34] builds upon traditional protein-ligand interaction repositories by curating data from the PDB with a specific emphasis on biological relevance. It compiles high-quality interactions, providing detailed annotations that help researchers better understand the functional roles of ligands in various protein environments. This makes BioLip a valuable resource for studying protein-ligand dynamics, particularly in drug discovery and enzyme catalysis. However, BioLip is restricted to experimentally validated data, limiting its scope to proteins structurally resolved through X-ray crystallography or NMR. This focus excludes proteins with predicted structures, which have become increasingly important with the rise of prediction technologies like AlphaFold.

With the advent of large-scale protein structure prediction, several new databases, such as HProteome-BSite [35], have emerged to catalog binding sites in AlphaFold-predicted structures. HProteome-BSite focuses on the human proteome, identifying potential ligand-binding sites across predicted protein models. This platform offers a broad view of human protein-ligand interactions, making it a valuable resource for drug discovery. However, it is limited to human proteins, leaving out essential insights for pathogens associated with neglected diseases.

CavitySpace [38] takes a more structural approach by identifying potential binding pockets in AlphaFold-predicted proteins. It provides detailed analyses of cavity sizes, shapes, and locations, crucial for understanding protein-ligand interactions and druggability. Though CavitySpace offers valuable geometric insights, its focus on human proteins and structural features needs to fully address the unique challenges of pathogen proteins, where different binding dynamics may occur.

AlphaFill [40] enhances AlphaFold-predicted models by integrating ligand information from experimentally solved structures in the Protein Data Bank (PDB). This database adds missing ligand-binding details, providing a more complete picture of predicted protein interactions. While AlphaFill enriches functional insights, it also primarily targets human proteins, limiting its utility in neglected diseases in which experimental data is sparse.

Current prediction tools have their strengths and weaknesses. Experimental data-driven databases like BindingDB and Binding MOAD offer high accuracy for known interactions but need more protein coverage with experimental data. On the other hand, databases using predicted structures provide broader coverage but may compromise prediction accuracy or specificity due to the inherent challenges in modeling binding sites based solely on computational methods. Moreover, few databases integrate multiple prediction tools, which limits their ability to provide a consensus view of binding sites, thereby introducing potential biases or inaccuracies inherent to individual algorithms.

In drug repurposing, several databases have been developed to assist in repositioning existing drugs by focusing on different aspects of drug-target interactions. DrugBank [42] and KEGG [43, 44] are foundational databases that offer basic but essential information about drugs and their targets, providing critical insights into biochemical pathways and interactions. However, these databases were not explicitly designed with drug repurposing in mind, often requiring researchers to extract relevant information manually, slowing the discovery process. Databases like RepoDB [45] and the Drug Repurposing Hub (DRH) [46] curate information about repurposed drugs, offering valuable data on both successful and unsuccessful repositioning efforts. While these resources play a crucial role in decision-making, they often need more sophisticated tools for proposing new drug applications.

Similarity-based databases such as PREDICT [47] and RepurposeDB [48] have emerged to address the need for more targeted repurposing tools. These platforms leverage molecular or target similarities to suggest new applications for existing drugs, but more than similarity is needed to fully capture the complexity of drug-target interactions. Additional biological, genetic, or clinical factors must be considered to improve the accuracy of these predictors. In addition, target-based databases like DMAP [49] and DrugSig [50] focus on identifying new drug applications by examining their interactions with multiple targets. Although effective in specific niche applications—such as orphan diseases or gene-specific therapies—these databases can be limited in scope and applicability. Moreover, the recently developed DrugR+ [51] addresses existing database limitations by integrating data from DrugBank and KEGG while offering enhanced functionalities. DrugR+ is designed to support both single and synthetic repositioning of drugs, providing researchers with new capabilities for predicting drug-target interactions using supervised machine learning methods.

DrugRepoBank [52] and the Therapeutic Target Database (TTD) [53] both offer comprehensive platforms that significantly support drug discovery and repositioning. DrugRepoBank integrates experimental data on repositioned drugs with predictive algorithms, including similarity-based, AI-driven, and network-based methods, to suggest new drug indications and targets. Its dual-module system combines literature-supported data with advanced predictive tools, facilitating both discovery and validation of drug repurposing efforts. Meanwhile, the TTD enriches drug discovery by cataloging comparative data on drugs and targets, such as poor binders, non-binders, and prodrug-drug pairs, while linking drug properties to the PDB and AlphaFold structural data.

While these drug repurposing databases and protein-ligand interaction catalogs are invaluable for advancing drug discovery, a significant gap exists in addressing diseases in which structural data is scarce, such as neglected diseases. This is where BENDER DB distinguishes itself by focusing specifically on binding site prediction for neglected disease pathogens using AlphaFold-predicted structures. Unlike existing databases that emphasize experimentally solved structures or focus on human proteins, BENDER DB is designed to catalog binding sites across multiple proteomes, addressing the limitations in current drug discovery pipelines by targeting pathogens that disproportionately affect vulnerable populations.

BENDER DB offers a comprehensive resource beyond individual techniques to provide consensus predictions by consolidating results from five state-of-the-art binding site prediction tools. This integration enhances the reliability of binding site identification, which is particularly crucial for neglected diseases in which accurate structural data is often unavailable. BENDER DB combines existing protein binding site databases and drug repurposing efforts, offering researchers a specialized platform to identify novel drug targets and provide actionable insights into therapeutic development.

Furthermore, BENDER DB’s interactive visual tools and user-friendly interface democratize access to high-quality binding site predictions, enabling researchers without extensive computational expertise to benefit from the platform and its data. This ease of access, coupled with its focus on neglected diseases, positions BENDER DB as a relevant resource for discovering and developing novel therapies in areas that traditional drug discovery approaches have historically underserved.

## 3. Methodology

BENDER DB involves four key components: data collection, binding site predictions calculated by five methods, the development of the metapredictor BENDER AI, and consensus-based visualization in the database. Data were collected from the proteomes of pathogens related to neglected diseases, with all protein structures obtained from the AlphaFold database. Binding sites were predicted using five state-of-the-art computational tools based on different paradigms to ensure comprehensive coverage. These results were then fed into BENDER AI, a meta-predictor designed to integrate the outputs of the various methods through a decision tree model. Finally, a consensus approach was employed to prioritize residues identified by multiple predictors, visualized in the BENDER DB database, providing an accessible platform for users to explore. The overall workflow is presented in Figure 1, which outlines the step-by-step methodology.

**Figure 1:**
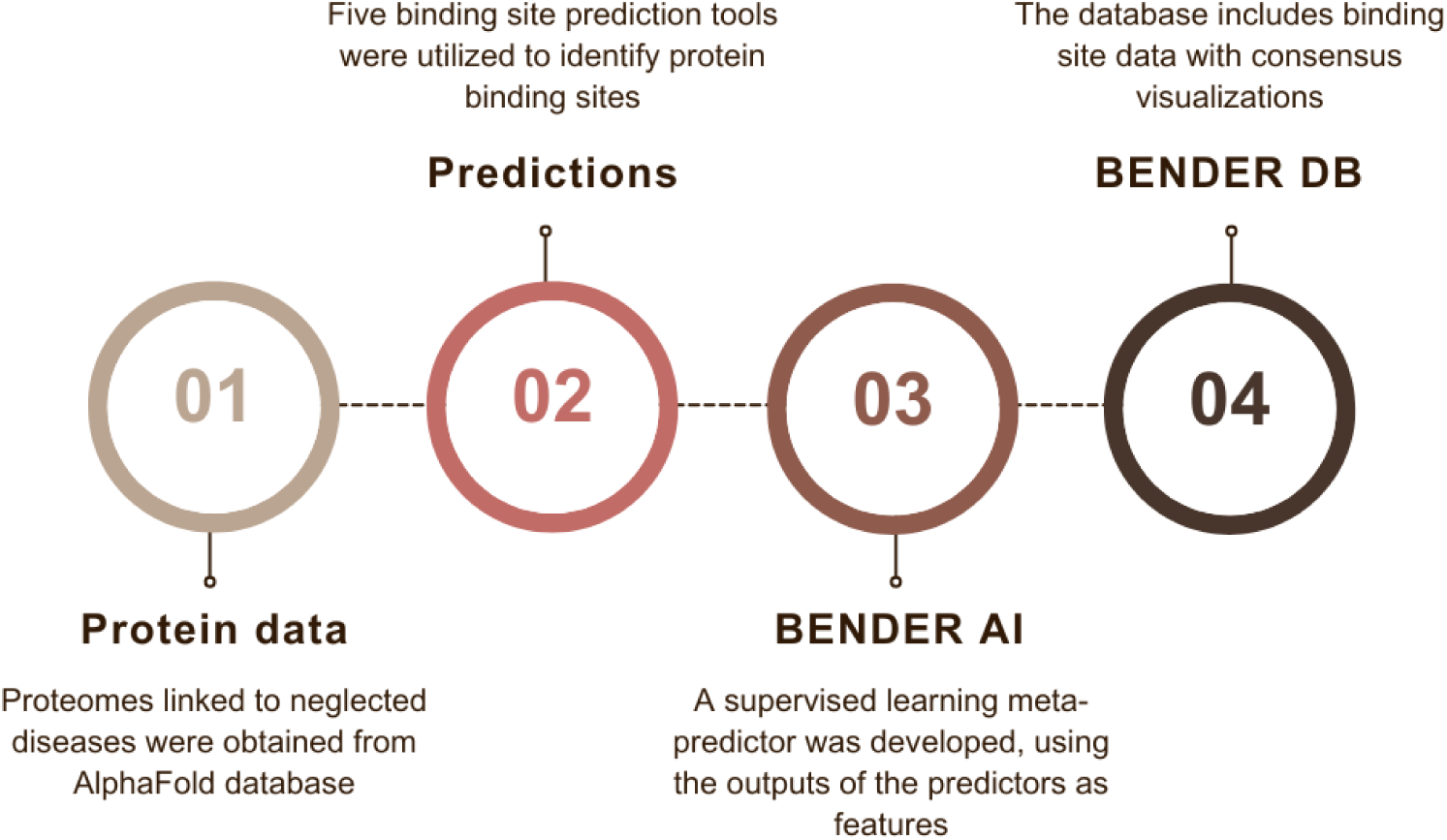
The workflow of BENDER DB is divided into four key components: (1) data collection from proteomes of pathogens with structures available in the AlphaFold database, (2) binding site prediction using five state-of-the-art computational methods, (3) meta-prediction performed by BENDER AI to aggregate results, and (4) visualization and exploration of binding sites through an interactive web interface.

### 3.1. Data collection

AlphaFold is a computational strategy to predict three-dimensional (3D) protein structures based on protein sequences. It performs structure prediction with atomic accuracy even when no similar structure is known. AlphaFold uses novel neural network architectures and training procedures based on protein structure’s evolutionary, physical, and geometric aspects. The data was collected from the AlphaFold Protein Structure Database, which stores all predicted structures made by the AlphaFold structure prediction method. The database is accessible at https://alphafold.ebi.ac.uk/ and conveniently organizes available data by proteomes, simplifying the extraction process.

Proteomes associated with pathogens causing neglected diseases, as listed by the World Health Organization (WHO) [54] and the Pan American Health Organization (PAHO) [55], were collected. Ten distinct proteomes were identified, resulting in 101,813 protein structures. Table 1 provides an overview of each proteome, including its name, the associated disease, and the number of proteins available on the AlphaFold database.

**Table 1:**
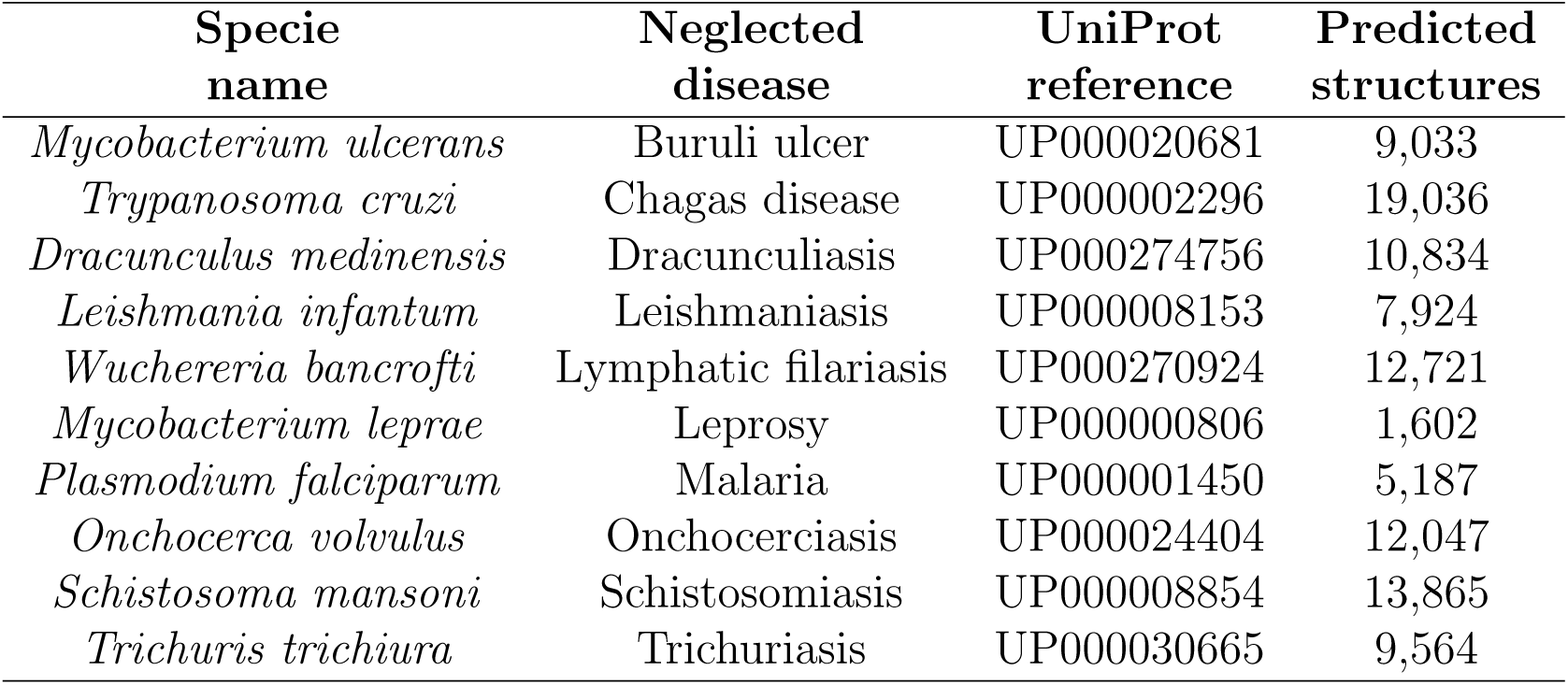
List of proteomes of neglected diseases pathogens obtained via AlphaFold database according to WHO and PAHO.

To ensure comprehensive coverage, we collected the complete proteome for each neglected disease pathogen in the AlphaFold database, using no exclusion criteria. We aimed to include all predicted structures to provide the most extensive resource possible for under-researched pathogens, which often lack sufficient scientific attention and resources. Although we recognize the potential inaccuracies inherent in AlphaFold predictions, we prioritized leveraging the entire dataset without additional preprocessing, such as alignment, filtering, or normalization. This approach facilitates large-scale exploratory analysis and enables the mapping of predicted binding sites across these proteomes, essential for advancing drug discovery in neglected diseases.

### 3.2. Binding site predictors

The binding sites of the collected proteins were calculated using five state-of-the-art computational strategies, each based on a distinct paradigm. The binding site predictors were selected according to the criteria: high performance reported in their recent publications, publicly available source code, and complementary approaches to binding site prediction (e.g., graph-based, machine learning, deep learning). This selection ensures the tools represent various methodologies, providing robust and diverse predictions. The outputs from these tools were standardized and integrated into BENDER DB by mapping all predictions to a standard residue format. Any potential discrepancies between the tools were addressed using a consensus-based approach, where the agreement among multiple predictors was prioritized to ensure higher confidence in the predicted binding sites.

GRaSP (Graph-based Residue neighborhood Strategy to Predict binding sites) introduces a supervised learning approach for predicting protein ligand-binding sites [56, 57]. GRaSP utilizes a residue-centric, graph-based strategy in which each residue and its neighbors are represented as a graph at the atomic level. Physicochemical and topological properties of atoms and non-covalent interactions are encoded as feature vectors, forming a matrix that serves as input for the supervised learning strategy. Unlike methods relying on sequence or structural alignment, GRaSP is scalable, residue-centric, and effective in identifying binding sites across multi-chain protein complexes.

The method calculates non-covalent interactions within a residue and its neighbor shells, summarized hierarchically. The graph model captures the physicochemical properties of the structural environment for each residue, represented as a matrix for supervised learning. The feature vector includes descriptors at residue, atom, and interaction levels. These descriptors encompass solvent accessibility, atom types, and various non-covalent interactions like hydrogen bonds and hydrophobic interactions. The binding site prediction problem is framed as a binary classification task, and GRaSP achieves compatible or superior results compared to state-of-the-art methods.

DeepPocket, a framework for efficient binding site detection in protein 3D structures, employs a multistep strategy [58]. Initially, candidate pocket centers are extracted using Fpocket, a geometry-based software known for accurately identifying pocket curvatures. Subsequently, a CNN-based classification model is employed for scoring these pockets, distinguishing positive samples based on their proximity to ligand atoms. To address data imbalance, the oversampling of positive examples is integrated. In the final step, a segmentation model inspired by the U-net architecture utilizes 3D CNNs to unveil pocket shapes at a precise resolution of 0.5 Å. Combining geometry-based tools, CNN scoring, and fine-grained segmentation, this integrated approach enhances the precision of binding site identification and ranking.

DeepPocket performed well, surpassing state-of-the-art methods at the time in identifying and ranking binding sites. The evaluation included diverse datasets and multiple benchmark criteria, underscoring the method efficacy during its assessment. With its modular design enabling independent utilization of components, DeepPocket proves to be a versatile tool for integration into structural bioinformatics and drug design pipelines. Focusing on fine-grained resolution contributes to heightened performance, which is particularly beneficial in scenarios where traditional template methods may fall short in providing essential binding site information.

PointSite [59] is a point cloud segmentation method designed to accurately identify protein ligand-binding sites (LBS) at the atom level. The approach begins by representing all atoms in a query protein as point clouds, labeling binding atoms with one and others with 0. The LBS identification problem is treated as a point cloud segmentation challenge, elegantly addressed by Submanifold Sparse Convolutional Networks (SSCN). Unlike traditional 3D convolutional neural networks (3D-CNN), SSCN avoids submanifold dilation issues and efficiently handles context learning by preserving sparsity in input points.

PointSite consists of three main modules: Atom Point Cloud Transformation (APCT), Ligand-binding Atoms Prediction (LAP), and LBS Identification (LBSI). The APCT module transforms the original protein structure into atom-level point clouds, labeling atoms based on their proximity to ligands. LAP predicts ligand atoms utilizing SSC-based U-net, enabling global context learning with sparse convolution operations. The LBSI module filters and reranks pocket-centric approach results based on the segmented ligand-binding atoms, offering a novel protein-centric perspective for LBS identification. The approach achieves state-of-the-art performance, especially in atom-level Intersection over Union (IoU) and Distance from the Center of Top-N identified LBS, even for challenging cases like unbound proteins.

PUResNet [60] is a deep-learning model that predicts ligand-binding sites on protein structures. The key innovation lies in developing an independent training dataset, a subset of scPDB, comprising 5,020 protein structures selected based on structural similarity, Tanimoto coefficient calculations, and manual inspection. PUResNet utilizes the ResNet architecture, consisting of encoder and decoder blocks with skip connections, to address the vanishing gradient problem commonly encountered in training deep neural networks. The protein structure is modeled as a 3D image (36 × 36 × 36 × 18), and PUResNet outputs a probability map (36 × 36 × 36 × 1) for each voxel, providing the likelihood of it belonging to a cavity.

In its architecture, PUResNet incorporates three basic blocks-convolution, identity, and up-sampling—inspired by both U-Net and ResNet concepts. The skip connections play a crucial role in mitigating the vanishing gradient problem. The model design emphasizes treating the protein structure as a 3D image, allowing it to predict ligand-binding sites effectively. PURes-Net architecture is described with specificity regarding the number of layers, parameters, and filter usage, offering a comprehensive understanding of its structure. The availability of the model and datasets on GitHub and the use of mol2 files for visualizing predictions ensures the reproducibility and practical application of PUResNet in predicting ligand-binding sites on protein structures.

P2Rank is a command line tool designed for predicting ligand binding sites in protein structures, offering a lightweight and platform-independent solution [61, 62]. Operating on Groovy and Java, the P2Rank binary package eliminates the need for additional dependencies and has been tested on both Linux and Windows systems. The tool efficiently processes input PDB files or datasets, producing automated predictions for each protein. The output includes CSV files containing detailed information about predicted pockets, including their coordinates, scores, solvent-exposed protein atoms, and constituent amino acid residues. Notably, the P2Rank algorithm classifies points evenly distributed on the protein Solvent Accessible Surface (SAS points), leveraging machine learning models to predict ligandability scores. The resulting high-scoring points are clustered to form predicted ligand binding sites, demonstrating the tool’s efficacy in identifying potential interaction sites.

The algorithm workflow involves generating regularly spaced SAS points, calculating feature descriptors based on local chemical neighborhoods, predicting ligandability scores through a Random Forest classifier, clustering high-scoring points to form pocket predictions, and ranking these pockets by cumulative ligandability scores. With 35 numerical features, the feature vector includes critical geometric aspects like protrusion, representing a point of buriedness. P2Rank efficiency is underscored by its quick runtime, processing a protein with 2,500 atoms in less than 1 second on a single core. This, coupled with its independence from external tools and databases, positions P2Rank as a valuable asset in structural bioinformatics pipelines, offering automated predictions and excelling in identifying novel allosteric sites.

Each predictor was used to identify binding sites in all collected proteins. The results were standardized and cataloged in the BENDER DB, providing a comprehensive resource for binding site analysis across multiple proteomes.

### 3.3. Binding site identification

The experiments were carried out using each method to identify binding sites. Protein-ligand binding sites were identified for the whole protein dataset, comprising all ten proteomes of neglected disease pathogens. Given a method, each of its binding sites was cataloged in the BENDER DB server. It is important to note that different predictors yield results in diverse formats.

GRaSP presents its results through a file containing a list of all residues belonging to the protein, each marked with a binary label indicating whether the residue belongs to a binding site. In the GRaSP version used, there is no residue clustering, so all results from GRaSP are considered a single binding site.

P2Rank results are organized based on the number of pockets detected. For each pocket, all residues belonging to that group are listed. Additional pocket characteristics, such as the number of surface atoms and the coordinates of the pocket center, are also included in the results.

PointSite outputs a list of all atoms in the protein, with each atom containing a label indicating the likelihood of belonging to a binding site. Residues with at least one atom labeled 0.5 or higher are considered part of a binding site.

DeepPocket often predicts many pockets in a given protein, but not all are high-confidence. For example, the results for protein Q9CD30 hold more than 100 pockets. To manage this, a 70% confidence threshold is used to determine whether a pocket is cataloged. Exceptions occur when no pockets exceed 70% confidence, in which case the five best-ranked pockets are noted, regardless of confidence.

PUResNet provides results in a list of pseudo-atoms and their coordinates, with no direct relationship to real atoms in the protein. To standardize results and represent PUResNet binding sites in residue format, residue mapping was performed based on the distance between predicted pseudo-atoms and the protein residues [63, 64, 65]. Pseudo-atoms within 5 Å of protein atoms were considered valid, and the corresponding residues were marked as belonging to the binding site.

BENDER DB brings together all this diverse information so that users can visualize and make sense of the binding sites and understand to what extent predictors agree. After calculating the binding sites for each protein of the neglected diseases proteomes, 1,172,743 binding sites were obtained and cataloged into the BENDER DB.

### 3.4. Data visualization

To present the results, BENDER DB employs two main visual representations: the NGL Viewer [66] for molecular visualization and the UpSet plot [41] for displaying intersections among methods results. In addition to listing the residues corresponding to a given binding site, visualizing these residues within the protein structure enriches the results, enabling users to perceive the spatial arrangement of the binding sites. For this purpose, we employed the NGL Viewer, an open-source web application designed to visualize, manipulate, and analyze molecular structures. Figure 2 shows the BENDER DB molecular viewer.

**Figure 2:**
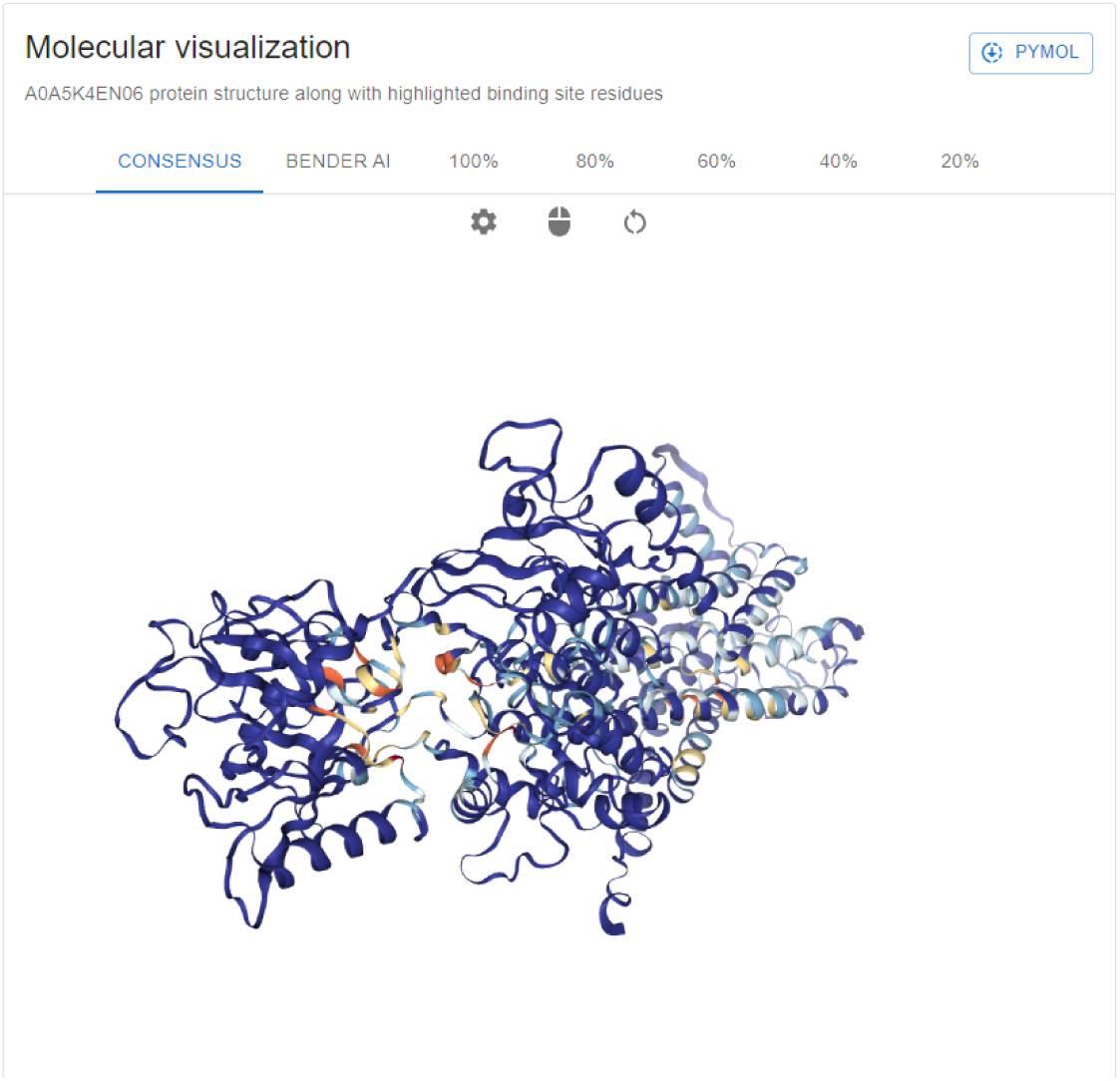
Molecular visualization page on BENDER DB. Tabs in the molecular viewer enable users to toggle between different graphical representations of the protein binding sites, providing an interactive platform for exploring residues and predictions.

When visualizing the results, the summary consensus tab presents a heatmap of all binding site residues. Darker shades of blue indicate a lower or no occurrence of binding site residues, as calculated by the predictors. In comparison, darker shades of red indicate a higher presence of binding site residues. The BENDER AI tab shows binding site residues predicted by our Artificial Intelligence model. The tabs with percentages show the occurrence of binding site residues calculated by all the predictors. For example, the 80% tab displays all binding site residues in at least 80% of the results.

The UpSet plot is a data visualization well-suited for displaying intersections among many sets. Figure 3 illustrates the key components of the UpSet plot:

- The lower-left area presents a list of all sets (in this study, corresponding to the results of the five binding site prediction methods). The horizontal bar size indicates the set cardinality, representing the total number of residues found in all binding sites for each predictor.
- The lower-right area showcases intersections between the sets. The lines in this area contain points that connect to indicate intersections between the predictors.
- The top area features vertical bars indicating the intersection’s cardinality. In our context, these vertical bars denote the total number of residues found in an intersection between different predictors.

**Figure 3:**
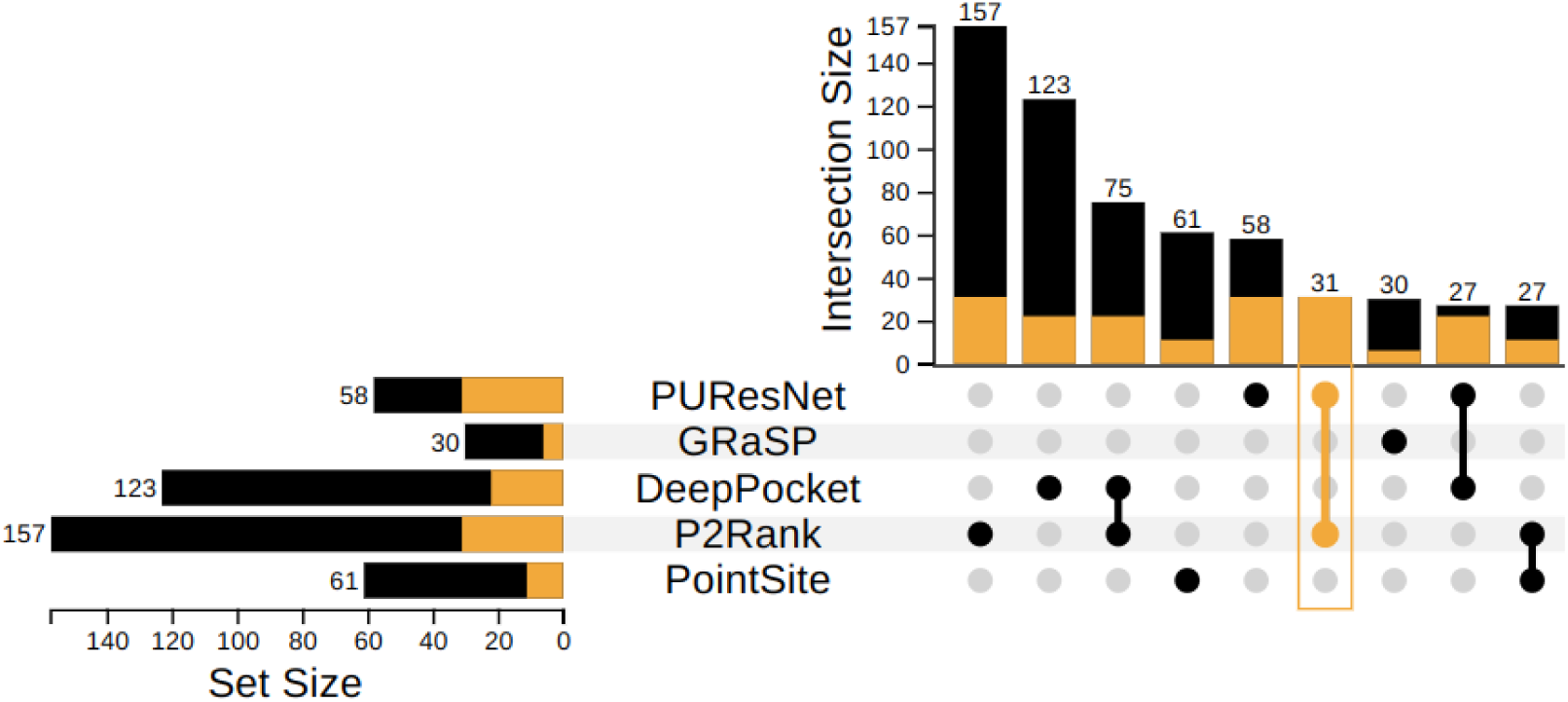
The UpSet plot displayed on the summary results page illustrates intersections and overlaps among binding site predictions from multiple tools. This visualization highlights agreement and divergence across the predictors, aiding in comparative analysis.

While the Venn Diagram is a common visual representation for illustrating intersections between sets, it becomes intricate and cluttered when dealing with four or more intersecting data sets. This is particularly relevant in the context of the BENDER DB, where there are five different sets (the binding site prediction results for each of the five binding site methods used). In the worst-case scenario, 31 intersections result in a confusing and congested Venn Diagram. Although only some of these intersections occur frequently, even having 15 or 20 intersections with five sets can make a Venn Diagram less effective. Thus, we opted for the UpSet plot, which enabled a more precise, cleaner, and focused visualization of methods results and their intersections. This representation facilitates gaining insights and conducting more precise analyses on points of agreement between the various predictors considered.

### 3.5. Web server implementation

To make the data of this work accessible, a web server has been developed, housing a database containing all the binding sites. BENDER DB was implemented using the Flask web framework (version 2.3.3) and the React library (version 18.2.0). The backend development, encompassing database analysis and processing, was undertaken using Flask. The front end, which includes all implemented pages and views, was designed using React. The web server is available at https://benderdb.ufv.br.

## 4. Results

### 4.1. Data report

A key feature BENDER DB offers is the ability to highlight how the binding side predictions of the various methods converge. While presenting the results of each predictor separately is important, indicating which residues are found in multiple predictors can offer valuable insights. A residue that consistently appears in the predictions of several predictors may carry greater significance than one found in only a single result. In response to this need, a summary tab has been created in the results section, providing a comprehensive overview of prediction results for each protein in the database.

Users can analyze the input protein on BENDER DB through two main views on the results page: the summary and individual predictor results. The summary view consolidates predictions into consensus results, providing an overview with three key areas: molecular visualization with a list of residues, predictors intersections, and binding site residue data, as shown in Figure 4. Users can further examine each prediction model separately in the individual predictor results view, with its molecular viewer for detailed, predictor-specific analysis.

**Figure 4:**
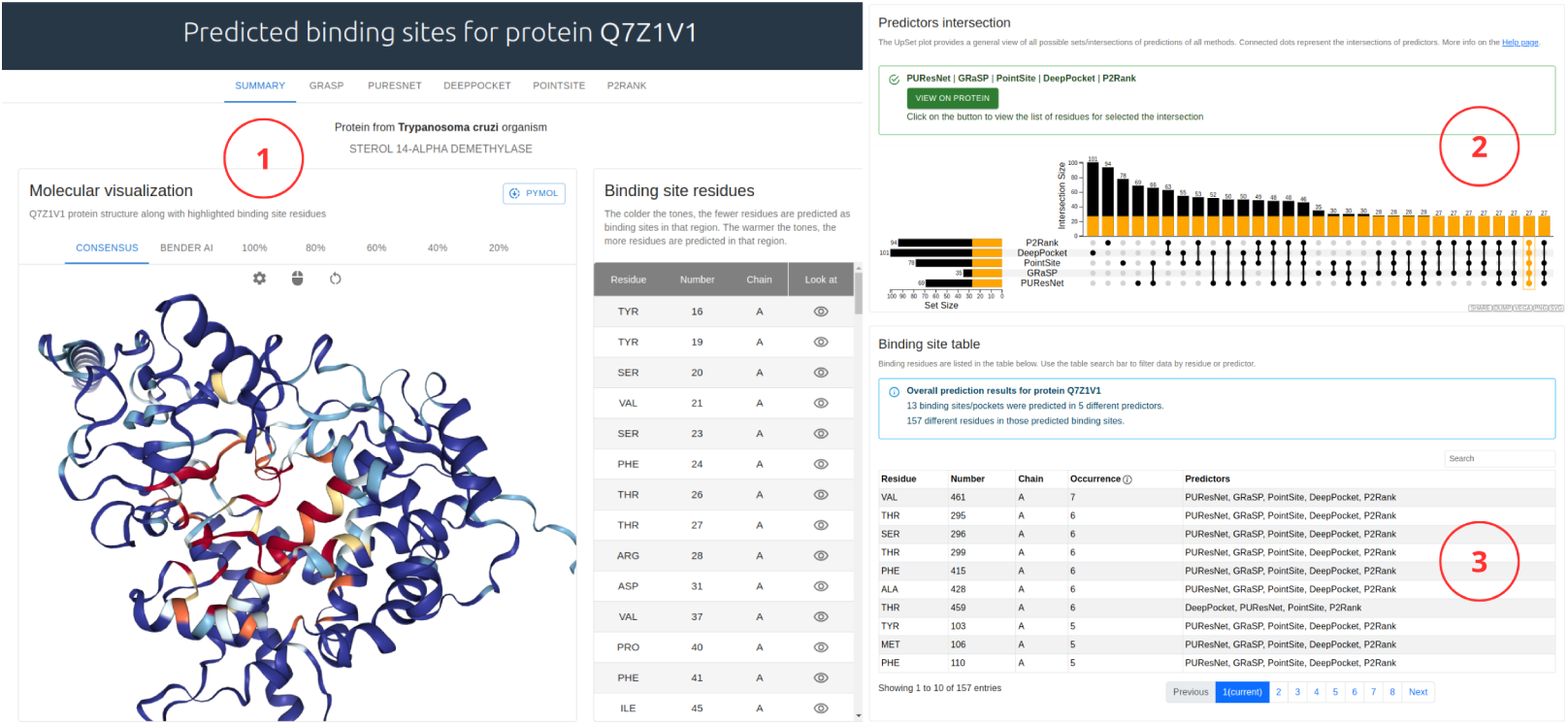
Overview of the results page on BENDER DB, displaying its three main sections. **(1) Molecular visualization:** The protein structure is shown alongside a table of residues, with tabs allowing users to toggle between different binding site visualizations. **(2) Predictors intersection:** The UpSet plot provides a comprehensive view of all possible sets and intersections of predictions across methods, with connected dots indicating predictor overlaps. **(3) Binding site table:** Binding residues are listed in a searchable table below, allowing users to filter data by residue or predictor for targeted analysis.

The molecular visualization offers various tabs displaying different representations of the protein. The consensus tab indicates the occurrence frequency of protein residues according to the predictions of the binding site identification models. This visualization employs a heatmap where cooler colors (blue tones) signify a low occurrence of the residue in prediction results, and warmer colors (red tones) indicate a higher occurrence. This enables users to identify residues and regions of the protein that are more likely to contain binding sites.

The BENDER AI tab employs a machine-learning model to classify protein residues as part of a binding site. This model utilizes features derived from the results of five binding site prediction methods, as discussed in Section 3.2.2. Residues identified as part of binding sites are highlighted in this visualization. Additionally, tabs show the percentage of residues considered binding sites according to the predictors. For instance, the 80% tab displays residues identified as binding sites by at least 80% (4 out of 5) of the predictors, regardless of which specific predictors they are. Next to the molecular visualization, a table listing residues is displayed, which updates according to the currently selected visualization tab.

The binding site intersections feature an Upset plot, a graphical visualization designed to analyze set intersections, as displayed in Figure 5. Each set represents the results of the binding site prediction methods. The UpSet plot is interactive and offers a variety of functionalities. First, users can click on objects on the graph, such as an intersection or data results from a particular predictor, to reveal relevant information. In the UpSet Plot itself, the bars (initially black, indicating totality) turn yellow, meaning that the selected intersection residues are present in other intersections and sets. Hovering the mouse over the sets and intersections before and after selection provides additional data regarding the total residues found. Upon selecting the subset of interest, users can visually analyze the protein structure using the molecular viewer by clicking the View on protein button (Figure 6).

**Figure 5:**
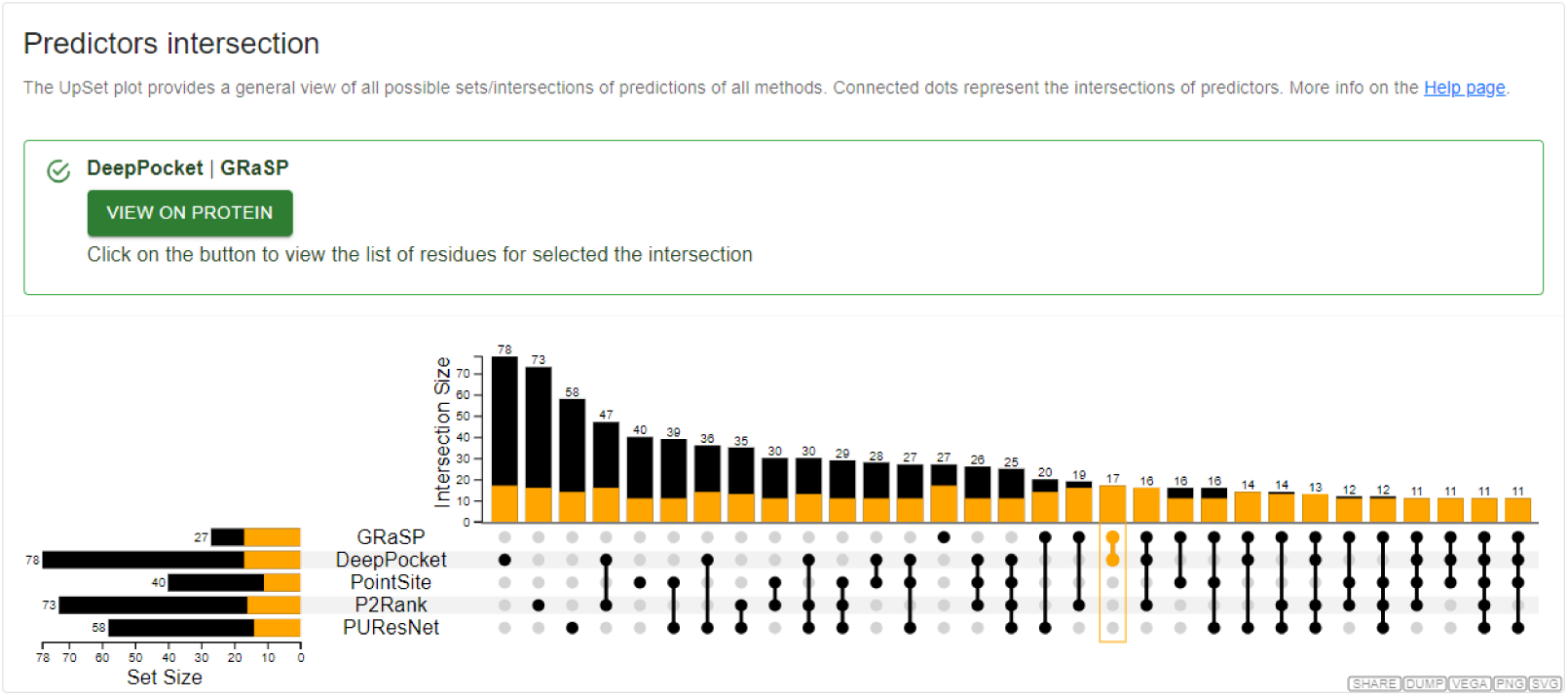
UpSet plot provides a comprehensive view of all tools’ sets and intersections of binding site predictions. This visualization highlights both agreements and discrepancies among prediction methods. The intersection of the GRaSP and DeepPocket predictors is selected, allowing users to see this subset’s total number of residues with more detail.

**Figure 6:**
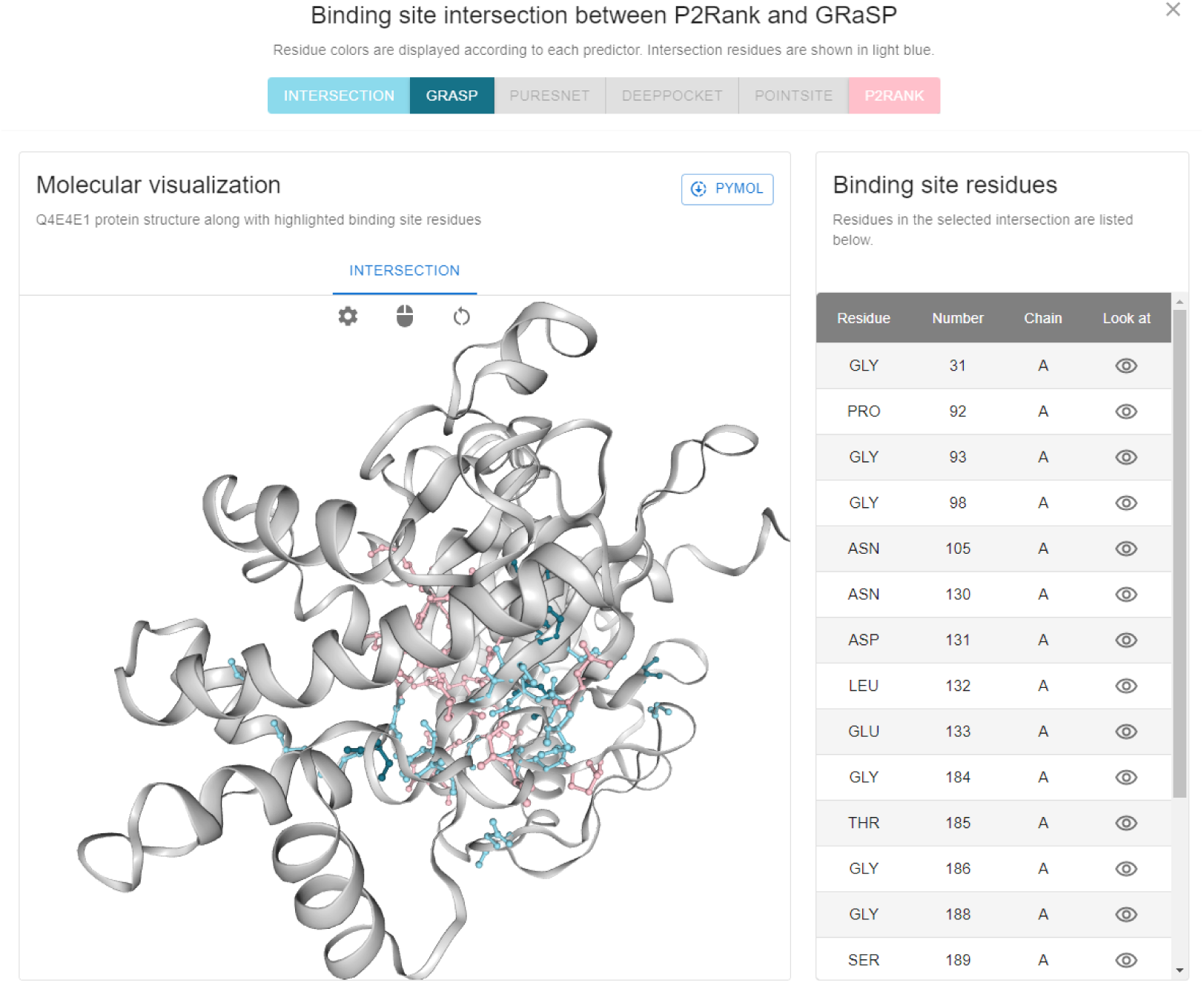
Pop-up window activated by clicking “View on protein” to analyze a selected intersection among predictor results. The molecular visualization highlights residues from the chosen intersection, displayed with colors assigned to each predictor. A list of the residues is also shown in the table on the right, enabling detailed exploration of the results.

The binding site residues data (Figure 7) provide information on the number of binding sites identified by the predictors, the number of residues forming these sites, and a table listing all cataloged residues and their occurrence in binding sites. While this overview of the results indicates the total number of residues and binding sites, a more in-depth analysis is necessary. This involves understanding which residues were most frequently found and examining each specific residue separately to discern which predictors did or did not identify a particular residue. An interactive table has been designed to address these aspects.

**Figure 7:**
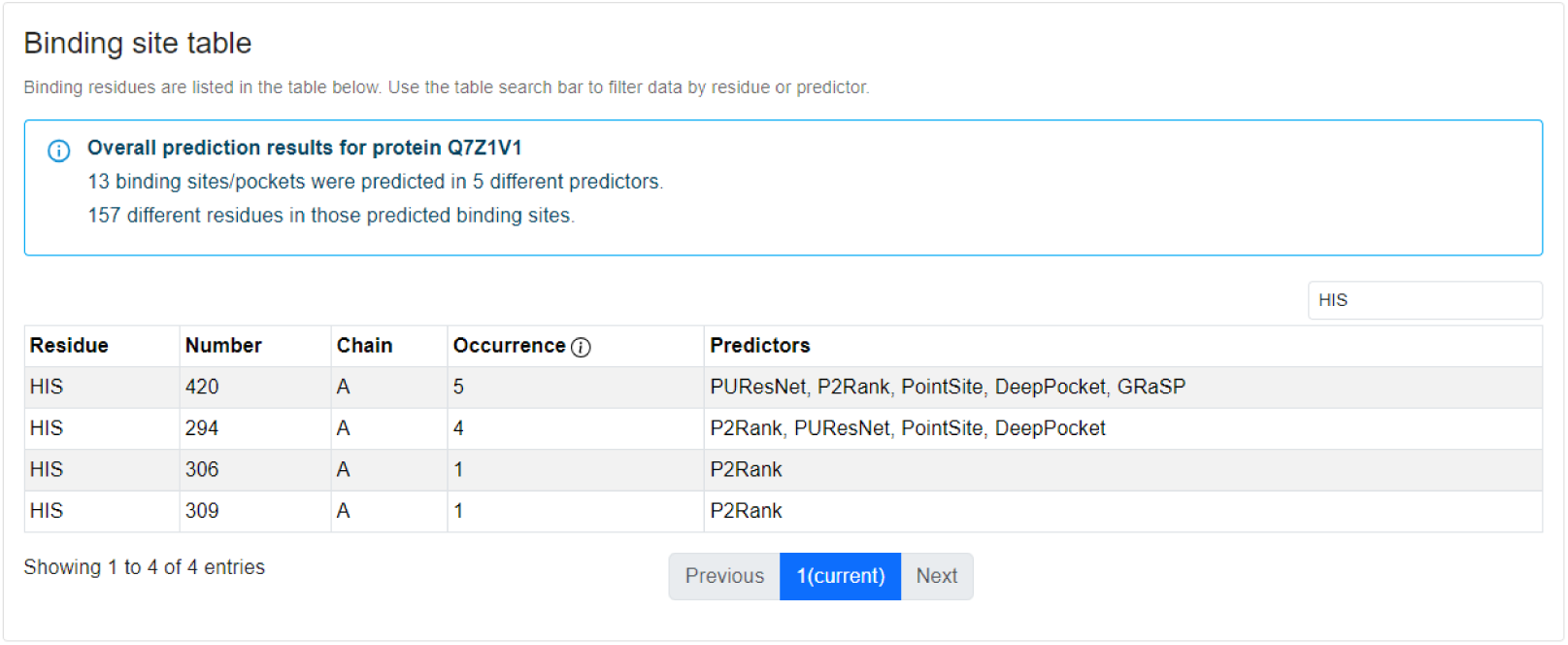
The binding site table is displayed on the summary results page. This interactive table lists the protein residues predicted as binding sites, allowing users to filter predictions by residue or prediction method for targeted exploration.

Beyond the summary tab, users can analyze the results of each predictor separately. The predictor tabs include a molecular visualization similar to the summary, allowing for detailed and specific analysis for each prediction model. Information regarding binding sites and their residues is presented within these tabs, accompanied by a molecule viewer illustrating these residues. Results can be downloaded as CSV files, encompassing the binding sites and their residues. It is also possible to download data from a specific method or a file containing data from all methods.

With these comprehensive tools, BENDER DB offers a robust and versatile platform for analyzing proteins and their binding sites. Those tools facilitate detailed and precise insights into the function and structure of the input protein, enhancing the understanding of this molecule.

### 4.2. BENDER consensus

Individual results from binding site predictions of the methods presented here allow the analysis of each case individually, according to the strengths of each method. However, these results often differ among the various predictors, making it difficult to determine whether residues can be considered part of the binding site.

The BENDER consensus was created to address this issue, aggregating all prediction results from various methods into a single representation. This allows for analyzing where methods converge and identifying residues more likely to belong to the protein binding site. Two summary representations of binding site residues are available: mean consensus and BENDER AI.

#### 4.2.1. Mean consensus

The consensus based on the mean of the predictions is calculated according to the percentage of occurrence of a residue in the results of all the methods. Each protein residue is categorized according to the percentage of predictors that consider it to belong to a binding site. The mean consensus groups the residues according to a probability threshold, and tabs show different agreement thresholds among the predictors. For instance, in the Tab 80% are presented residues detected as part of a binding site by at least 80% of the predictors. Figure 8 highlights the various tabs with the occurrence percentages. When one of these tabs is selected, the table on the right-hand side of the molecular viewer updates to show the residues listed by the selected tab. An extra representation for this consensus was created, grouping all occurrence percentages into a single visualization, the consensus tab, as described in Section 3.4.

**Figure 8:**
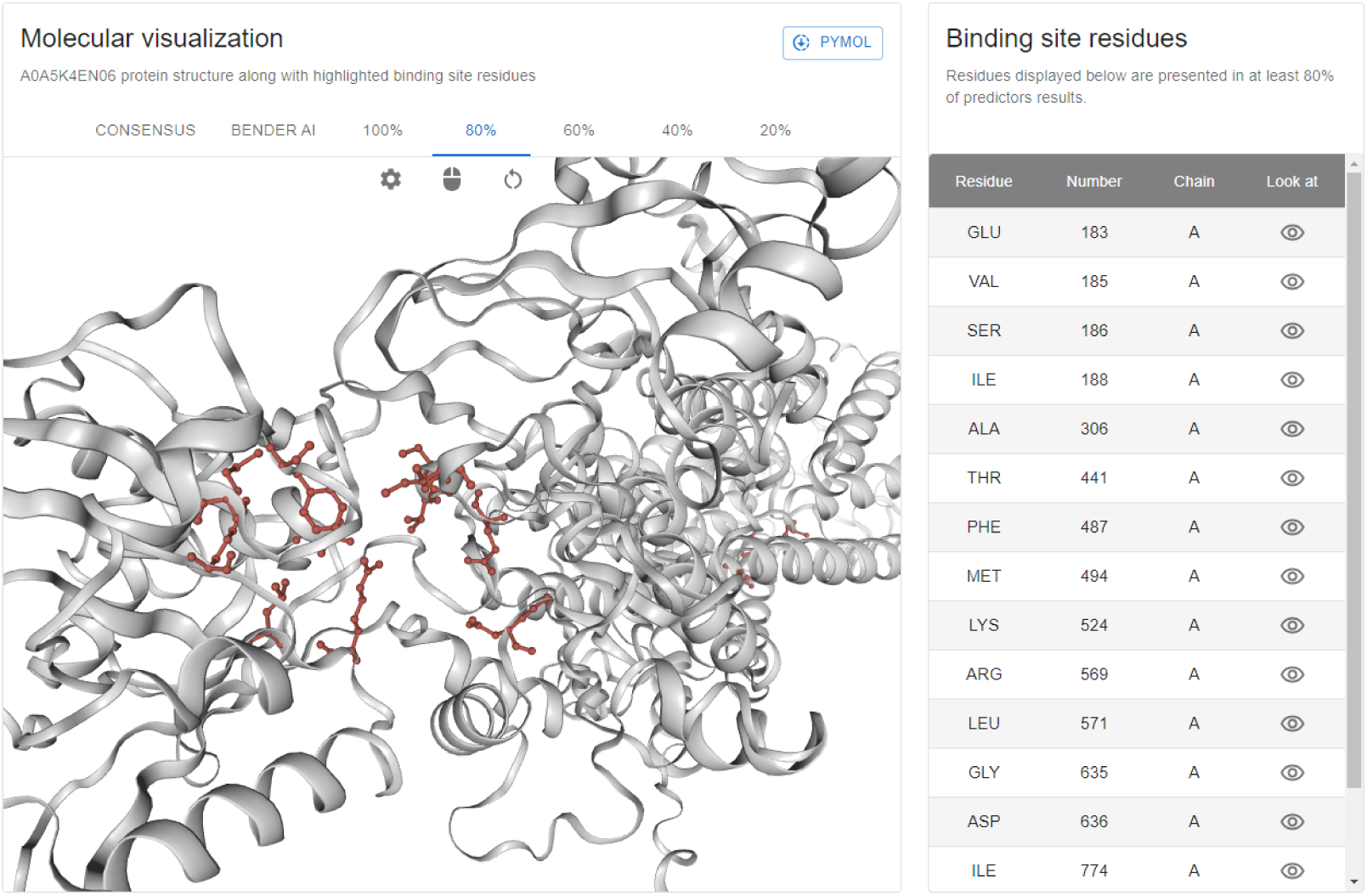
Structural representation of the protein shown alongside a table listing residues identified as binding sites. These are consensus results, displaying residues at least 80% of the predictors identify as binding sites. The table on the right lists these residues, and users can switch between tabs to analyze different levels of predictor convergence.

#### 4.2.2. BENDER AI

To provide an additional perspective on residues belonging to binding sites, we developed BENDER AI, a supervised learning strategy designed to identify binding sites in proteins by integrating the outputs of several established predictors. BENDER AI is a meta-predictor incorporating the results of the five predictors used in this work, using their outputs as features. Each residue in a protein is characterized by a binary value: 1 if a predictor identifies it as part of a binding site and 0 otherwise. This integration allows BENDER AI to combine the predictions into a cohesive classification model.

For the training and testing of BENDER AI, we used datasets provided by COACH [67]. The training dataset consisted of 400 non-redundant proteins, while the test dataset comprised 500 non-redundant proteins containing 814 ligands. We employed TPOT, a Python tool that automates machine learning pipeline optimization through genetic programming, to determine the most effective machine learning model [68]. TPOT identified the decision tree classifier as the optimal model, which was subsequently used for our classification tasks. The decision tree was configured with the following parameters: the **criterion** was set to “entropy”, which allows the model to measure the quality of a split using information gain; the **max depth** was limited to 4 to prevent overfitting; and both **min samples leaf** and **min samples split** were set to 16 to ensure that splits occurred only when there were a sufficient number of samples, promoting generalizability. These parameters were optimized based on the average cross-validation accuracy score during training, which achieved 0.9633. Following the creation of the model, we performed predictions on all 101,823 proteins across the proteomes covered by BENDER DB. These results are available in the web server results page summary under the BENDER AI tab.

We used several well-established classification metrics to evaluate the performance of BENDER AI in predicting protein binding sites: Matthews Correlation Coefficient (MCC), Precision, Recall, F1, and AUC scores. Each of these metrics was chosen for its relevance in binding site prediction, providing a comprehensive assessment of the model’s ability to make accurate and reliable predictions.

MCC is a robust measure that considers true and false positives and negatives, providing a balanced metric even for imbalanced datasets. In the context of protein binding site prediction, MCC is particularly important as it evaluates the overall reliability of the predictions, accounting for both correct and incorrect classifications. This is essential in imbalanced data. In the binding site prediction problem, the number of residues that are part of binding sites is significantly smaller than that of non-binding residues.

Precision measures the proportion of predicted binding sites that are actual binding sites. In drug discovery, high precision is crucial as it minimizes the number of false positives, ensuring that the identified binding sites are likely to be actual binding sites. This is especially important when the model guides experimental validation, in which false positives could lead to wasted time and resources.

Recall (or sensitivity) evaluates how many of the actual binding site residues were correctly recovered by the model. In neglected diseases, recall is critical because it ensures that potential drug targets are not missed. A higher recall indicates that the model captures all possible binding sites more effectively, which is important for comprehensive drug discovery efforts. The F1 score is the harmonic mean of precision and recall, balancing these two metrics trade-offs. In protein binding site prediction, the F1 score is helpful because it provides a single metric that considers both the need to minimize false positives (precision) and identify as many actual binding sites as possible (recall). This balance is crucial when designing experiments based on the model predictions.

The AUC (Area Under the Curve) score measures the model’s ability to differentiate between binding and non-binding residues across various thresholds. A high AUC indicates strong classification performance in protein binding site prediction, showing that the model consistently ranks true binding sites above non-binding residues. This makes AUC particularly valuable in imbalanced datasets, providing a reliable assessment of overall prediction quality.

These metrics are important in protein binding site prediction, in which imbalanced datasets are common. The number of non-binding residues often far exceeds the number of actual binding site residues, making accuracy an insufficient performance measure. MCC, precision, recall, F1 score, and AUC offer a more nuanced assessment, each addressing different aspects of model performance.

By evaluating BENDER AI with these metrics, we ensure a thorough understanding of the strengths and weaknesses of the model. This approach provides a well-rounded evaluation, making it possible to assess both the reliability of the predictions and the model’s ability to capture relevant binding sites. In neglected diseases, identifying viable drug targets is critical. These metrics help evaluate the model’s effectiveness and ensure it can contribute meaningfully to drug discovery efforts.

The evaluation results are presented in Table 2. BENDER AI demonstrates strong performance across multiple evaluation metrics, achieving the highest MCC, precision, F1, and AUC scores among all the compared tools. The AUC score, reaching 0.89, highlights BENDER AI’s ability to distinguish true binding sites consistently, reinforcing its predictive strength. These metrics reflect the accurate predictions of our method, highlighting its capacity to distinguish actual binding sites while maintaining a balance between precision and recall. The high precision score and the highest AUC indicate that BENDER AI effectively minimizes false positives, making it a reliable predictor for binding site identification.

**Table 2:** Evaluation metric scores achieved by BENDER AI compared to other predictors using the test dataset.

| Metric | BENDER AI | GRaSP | PUResNet | DeepPocket | PointSite | P2Rank |
| --- | --- | --- | --- | --- | --- | --- |
| MCC | **0.64** | 0.63 | 0.33 | 0.31 | 0.44 | 0.33 |
| Precision | **0.70** | 0.62 | 0.23 | 0.20 | 0.30 | 0.22 |
| Recall | 0.61 | 0.67 | 0.65 | 0.72 | **0.77** | 0.66 |
| F1 score | **0.65** | **0.65** | 0.34 | 0.31 | 0.43 | 0.33 |
| AUC | **0.89** | 0.82 | 0.76 | 0.78 | 0.84 | 0.77 |

However, while BENDER AI excels in precision and AUC, its slightly lower recall score than other predictors suggests it may miss some actual binding sites. This trade-off is common in predictive models in which high confidence is placed on positive predictions, potentially at the cost of identifying all true positives. To address the lower recall, future improvements could include adjusting the decision thresholds to increase sensitivity or incorporating more diverse training data to better capture different binding sites. Additionally, combining predictions from multiple models could further enhance recall by expanding coverage. Despite the current trade-off, the overall balanced performance across MCC, precision, F1, and AUC scores confirms BENDER AI’s robust capability to predict binding sites with significant reliability. These strengths underscore its protein binding site prediction value, particularly in providing high-confidence results.

### 4.3. Case study

Protein binding site identification plays a critical role in drug discovery and repositioning, as it provides essential information about where potential small molecules or drugs can interact with proteins [69, 70]. By predicting and validating binding sites, researchers can identify new drug targets, optimize drug designs, or repurpose existing drugs for new therapeutic uses [11, 71]. In recent years, advances in computational techniques, such as machine learning, molecular docking, and deep learning, have significantly improved the accuracy and efficiency of binding site predictions, making them valuable tools in early-stage drug discovery [72, 73, 74, 75]. For instance, modern prediction tools allow for the identification of binding pockets that were previously overlooked, facilitating the design of more effective inhibitors and modulators [76].

Drug repositioning benefits from these computational techniques, enabling existing compounds to target newly identified binding sites and reducing development costs and timelines [77]. Binding site predictions have been successfully applied to identify alternative uses for approved drugs by matching them to similar binding sites in different proteins, which has led to several repurposed therapeutics [78, 79]. For example, predictions identified Oseltamivir, Etravirine, and Fosamprenavir as high-affinity inhibitors of SARS-CoV-2 key proteins, such as the Spike-ACE2 complex, highlighting their potential as repurposed COVID-19 therapeutics [80].

These approaches are especially advantageous for neglected diseases, in which resources for traditional drug discovery are limited. Computational methods provide a cost-effective way of exploring therapeutic options and accelerating drug discovery. In this context, BENDER DB offers a platform that integrates binding site predictions for proteins from neglected diseases, supporting further research into drug candidates and repurposing opportunities.

To illustrate how results from BENDER DB can be applied in drug discovery and repositioning studies, we focus on Sterol 14-alpha demethylase (CYP51) with the UniProt accession Q7Z1V1. This enzyme is critical in the biosynthesis of ergosterol, an essential component of cell membranes [49]. CYP51 belongs to the cytochrome P450 superfamily, characterized by a conserved P450 protein fold that includes alpha-helices and beta-sheets. Due to its role in sterol biosynthesis, the enzyme is vital in various organisms, including fungi, parasites, and humans. In *Trypanosoma cruzi*, the parasitic organism responsible for Chagas disease, CYP51 is essential for the survival and proliferation of the organism.

Given its essential role, CYP51 is a prime target for antifungal drugs such as azoles. These drugs inhibit the enzyme activity, disrupting membrane integrity and leading to the death of the target organism. The active site of CYP51 is designed to bind substrates and inhibitors, allowing for precise interactions that influence enzyme activity. Understanding these interactions is crucial, as mutations in the CYP51 gene can lead to drug resistance, particularly in fungi and parasites.

The Protein Data Bank (PDB) contains 21 cataloged structures of CYP51, offering valuable insights into its three-dimensional conformation. These structures aid in understanding the molecular mechanisms of drug action and resistance, facilitating the development of new therapeutics with improved efficacy and minimized side effects. The detailed study of CYP51 structures, especially in complex with antifungal drugs, provides a foundation for designing novel treatments against fungal and parasitic infections, ultimately contributing to better managing diseases caused by these pathogens.

One protein in the polymer entity group content that matches the UniProt accession Q7Z1V1 is the protein with PDB code 2WUZ, which has 25 residues cataloged as binding sites in the PDB and 18 in BioLip. Both BENDER AI and the BENDER consensus successfully identified these binding residues. BENDER AI correctly identified 22 out of the 25 residues cataloged by the PDB and 17 out of the 18 cataloged by BioLip, demonstrating its ability to accurately predict binding sites with minimal false positives. Similarly, the BENDER consensus results further supported this accuracy, especially in the **100% consensus tab** (residues identified by all predictors), where 22 residues matched the PDB annotations and 17 matched BioLip. Expanding to the **80% consensus tab** (residues identified by at least four predictors), the consensus identified 24 of the 25 PDB residues and all 18 BioLip residues. We compared BENDER AI with the other predictors to numerically analyze performance using the precision metric. The results presented in Table 3 highlight the precision of the predictors in identifying protein binding sites for UniProt accession Q7Z1V1 (PDB code 2WUZ). Although all predictors successfully identified the binding site residues cataloged in PDB and BioLip, the main issue lies in the high number of false positives produced by most methods, significantly impacting their precision values. For instance, DeepPocket identified 24 residues correctly according to PDB but predicted 101 residues, resulting in a low precision of 0.24. Similarly, other methods, such as PUResNet and PointSite, struggled with high false positive rates, yielding precision values of 0.36 and 0.37 for PDB, respectively. In contrast, BENDER AI demonstrated a clear advantage, achieving the highest precision values of 0.79 for PDB and 0.61 for BioLip. These results emphasize BENDER AI’s robustness in minimizing false positives and providing reliable binding site predictions, making it the most effective tool among the evaluated predictors.

**Table 3:** Comparison of precision values for different predictors in identifying protein binding sites on UniProt accession Q7Z1V1, corresponding to the protein with PDB code 2WUZ. The columns represent the following: **Predictor** refers to the prediction model being evaluated; **TP PDB** is the number of true positives (binding residues) correctly identified based on PDB annotations; **TP BioLip** is the number of true positives correctly identified based on BioLip annotations; **Total Predicted** is the total number of residues predicted as binding sites by each predictor; **Precision PDB** is the proportion of correct predictions (true positives) out of all predictions according to PDB; **Precision BioLip** is the same proportion based on BioLip annotations.

| Predictor | TP PDB | TP BioLip | Total Predicted | Precision PDB | Precision BioLip |
| --- | --- | --- | --- | --- | --- |
| GRaSP | 22 | 17 | 35 | 0.63 | 0.49 |
| PUResNet | 25 | 18 | 69 | 0.36 | 0.26 |
| DeepPocket | 24 | 18 | 101 | 0.24 | 0.18 |
| PointSite | 25 | 18 | 78 | 0.37 | 0.26 |
| P2Rank | 23 | 17 | 94 | 0.24 | 0.18 |
| BENDER AI | 22 | 17 | 28 | **0.79** | **0.61** |

With these comprehensive tools, BENDER DB offers a robust and versatile platform for analyzing proteins and their binding sites. This facilitates detailed and precise insights into the function and structure of proteins from neglected disease pathogens organisms, such as the Sterol 14-alpha demethylase (CYP51) protein.

## 5. Limitations and future work

While BENDER DB is a valuable resource for studying protein binding sites, some limitations should be noted. The database relies on AlphaFold-predicted structures, which are still based on computational models despite being highly accurate and widely regarded as the standard in protein structure prediction. These predictions do not always fully capture the complexity of biological environments, especially in cases where protein interactions and dynamics play a significant role. Thus, while AlphaFold provides a strong foundation, users should remember that some predictions may not wholly reflect real biological contexts.

BENDER DB does not include experimental validation of the predicted binding sites. While computational predictions offer significant advantages in speed and cost, experimental validation is essential for confirming the accuracy of these predictions. However, incorporating such validation for every predicted binding site would require extensive resources and is outside the scope of this work. In future work, we aim to collaborate with experimental biologists to validate a subset of predictions, which would further strengthen the reliability of the database.

Currently, BENDER DB operates with pre-calculated results and does not allow users to input new protein sequences for analysis. This static nature limits the database’s flexibility in accommodating new research needs. Future improvements will focus on expanding user interactivity, enabling researchers to analyze new proteins and binding sites. This enhancement will allow for comparisons with existing data in the database, increasing its relevance and utility in ongoing studies and broadening the platform’s impact.

There are many directions for expanding BENDER DB capabilities. Including additional proteomes, particularly those related to neglected diseases, would broaden the scope of the proposed tool. Integrating more binding site prediction methods could also enhance the robustness of the predictions, offering a more comprehensive analysis. Furthermore, introducing features like real-time updates and more advanced querying options would improve user experience and increase the platform’s adaptability.

## 6. Conclusion

In this work, we developed BENDER DB, a protein binding site database across neglected disease pathogens’ proteomes. By integrating the predictions from five state-of-the-art binding site methods, BENDER DB offers a comprehensive and robust resource that can aid researchers in identifying viable drug targets more efficiently. The database uses structures predicted by AlphaFold and includes interactive visual tools, which provide a detailed and practical platform for supporting drug discovery and research in neglected diseases.

Integrating multiple predictors in BENDER DB highlights the convergences and differences among various prediction methods and offers a detailed analysis through tools like the UpSet plot and molecular visualizations. The addition of BENDER AI, a supervised learning strategy that combines the prediction of five methods, further enhances the accuracy and reliability of binding site identification. By aggregating predictions into a single representation, BENDER AI helps to resolve discrepancies among individual predictors, offering a consensus that can guide researchers in pinpointing the most promising binding sites.

Ultimately, BENDER DB’s comprehensive mapping and advanced analytical tools provide an invaluable resource for the scientific community. This platform can accelerate drug discovery efforts for neglected diseases by providing accurate and detailed information on protein binding sites. Through its innovative features and user-friendly interface, BENDER DB stands to significantly impact the identification and development of effective treatments, thereby addressing the urgent need for better therapeutic options in underdeveloped and developing regions.

## Acknowledgements

Funding: This work has been supported by funding from the Coordenação de Aperfeiçoamento de Pessoal de Nível Superior (CAPES), Conselho Nacional de Desenvolvimento Científico e Tecnológico (CNPq), Fundação de Amparo à Pesquisa do Estado de Minas Gerais (FAPEMIG).

## Declaration of generative AI and AI-assisted technologies in the writing process

During the preparation of this work, the author(s) used Grammarly and ChatGPT in order to improve language, readability, and perform grammar checks. After using these tools, the author(s) reviewed and edited the content as needed and take(s) full responsibility for the content of the publication.

